# Sensorimotor Memory for Object Weight is Based on Previous Experience During Lifting, Not Holding

**DOI:** 10.1101/464693

**Authors:** Vonne van Polanen, Marco Davare

## Abstract

To allow skilled object manipulation, the brain must generate a motor command specifically tailored to the object properties. For instance, in object lifting, the forces applied by the fingertips must be scaled to the object’s weight. When lifting a series of objects, forces are usually scaled according to recent experience from previously lifted objects, an effect often referred to as sensorimotor memory. In this study, we investigated the specific time period during which stored information from previous object manipulation is used to mediate sensorimotor memory. More specifically, we examined whether sensorimotor memory was based on weight information obtained between object contact and lift completion (lifting phase) or during stable holding (holding phase). Participants lifted objects in virtual reality that could increase or decrease in weight after the object was lifted and held in the air. In this way, we could distinguish whether the force planning in the next lift was scaled depending on weight information gathered from either the dynamic lifting or static holding period. We found that force planning was based on the previous object weight experienced during the lifting, but not holding, phase. This suggest that the lifting phase, while merely lasting a few hundred milliseconds, is a key time period for building up internal object representations used for planning future hand-object interactions.

**HIGHLIGHTS:**

- When lifting objects, fingertip force scaling is based on the most recent lift
- We investigated what time period is critical for acquiring sensorimotor memory
- Sensorimotor memory is based on weight experienced during previous lift, not hold
- The lifting phase is a key period for building up internal models of object lifting

## 1 INTRODUCTION

Humans have the exquisite capability to use their hands and manipulate objects skilfully with effortless grace. Skilled object manipulation requires the brain to issue a motor command specifically tailored to the physical object properties. For instance, when lifting an object, the forces applied by the fingertips must be scaled to the object’s weight. Object weight cannot be directly perceived before the object is lifted, but it can be estimated from other object properties such as its material, size (Gordon, Forssberg, Johansson, & Westling, 1991), and even learned arbitrary cues (Ameli, Dafotakis, Fink, & Nowak, 2008; Chouinard, Leonard, & Paus, 2005) in order to appropriately plan fingertip forces in a feedforward manner. If forces are planned incorrectly, for instance when anticipating a heavy object that is in fact light, they are quickly adjusted during the lift based on rapid sensory feedback loops (Johansson & Flanagan, 2009; Johansson & Westling, 1984, 1988a). Forces can also be quickly adapted if they are perturbed during the stationary holding of an object (Cole & Abbs, 1988; Johansson & Westling, 1988b) or even during lifting (Mrotek, Hart, Schot, & Fennigkoh, 2004).

When information about object weight cannot be inferred from viewing the object, such as in lifting a series of similarly looking objects that have different weights, force planning is based on the most recent lift. This phenomenon is known as sensorimotor memory (Johansson & Westling, 1988a; Loh, Kirsch, Rothwell, Lemon, & Davare, 2010; van Polanen & Davare, 2015). For example, if a lifted object was preceded by the lift of a heavy object, the planned forces are higher than when a light object was lifted previously. Sensorimotor memory for objects can last for hours (Flanagan, King, Wolpert, & Johansson, 2001; Green, Grierson, Dubrowski, & Carnahan, 2010; Nowak, Koupan, & Hermsdorfer, 2007) and is represented in the primary motor cortex (Chouinard, et al., 2005; Loh, et al., 2010).

Although sensorimotor memory and force scaling for objects has been a well-investigated topic over the past decades, it remains unclear what kind of information it is based on. Some studies suggest that it represents the memory of a force or is related to the sensation of a force. Evidence for this comes from studies that found a disruption of sensorimotor memory by wrist angulation (Bensmail, Sarfeld, Fink, & Nowak, 2010) or isometric forces (Quaney, Rotella, Peterson, & Cole, 2003). Furthermore, vibrations applied to the hand disrupt sensory signals and this was also shown to affect sensorimotor memory (Nowak, Rosenkranz, Hermsdorfer, & Rothwell, 2004). These studies indicate that sensorimotor memory consists of a memory of the force that is last performed and is not necessarily related to object properties.

On the other hand, another experiment contradicts these results by failing to disrupt sensorimotor memory with an isometric force task (Cole, Potash, & Peterson, 2008). Furthermore, there is evidence that sensorimotor memory for objects can be transferred between hands (Chang, Flanagan, & Goodale, 2008; Gordon, Forssberg, & Iwasaki, 1994; Green, et al., 2010) and there is also some transfer across tasks (Parikh & Cole, 2011). It has also been suggested that sensorimotor memory represents a memory of visual object size, as participants adjusted their forces correctly to a slightly smaller object without consciously noting the size change (Cole, 2008). Another example of force scaling towards new objects based on previous experience is the extrapolation of forces when lifting a sequence of increasing weights. Here, force scaling was not based on the previous lift, but forces were extrapolated based on the previous series (Mawase & Karniel, 2010). This literature suggests that sensorimotor memory is a representation of an object, or an internal model about object properties, that can be retrieved to plan forces when lifting a similar object.

An issue that has been unaddressed until now is *when* sensorimotor memory for an object is acquired. A lifting movement can be divided into loading phase (from object contact until lift-off), lift, hold (when the object is held stable in the air), replacement and unloading (from object contact with the table surface until release of the fingers from the object) (Johansson & Flanagan, 2009). The weight of an object can already be estimated at the moment of lift-off, i.e. when fingertip forces overcome object weight and the object starts to move. At this time point, corrective feedback processes are important to adjust forces and ensure a stable holding of the object. Here, we will refer to the period from object contact until the object reaches the target lifting height in the air as the ‘lifting phase’ and the phase when the object is held still in the air at the target height as the ‘holding phase’.

The lifting phase has been found to be important to mediate perception of weight when actively lifting objects (van Polanen & Davare, 2015) or from observing others lifting objects (Hamilton, Joyce, Flanagan, Frith, & Wolpert, 2007). This could indicate that this phase is important for establishing object weight and creating a sensorimotor memory for the object. However, it must be noted here that weight perception is not always related to force scaling. For example, in the size-weight illusion where two differently sized but equally weighting objects are lifted, the objects are perceived as having different weights whereas they are lifted with similar forces (Chang, et al., 2008; Flanagan & Beltzner, 2000; Grandy & Westwood, 2006).

In contrast to the hypothesis that the lifting phase is most important for building up sensorimotor memory, the holding phase could also provide important information. The weight that is perceived during steady holding and release of the object will be the latest sensory input about the object that is manipulated. Since sensorimotor memory is generally based on the most recent handled object, it is plausible that this final sensory information could be used to plan the next lift. Furthermore, the stable holding phase allows collection of sensory inputs over a much longer time period than the lifting phase, without movement-induced sensory noise, altogether providing more accurate information about weight, hence building a more reliable sensorimotor memory.

To investigate which time period, i.e. lifting vs. holding phase, sensorimotor memory is based on, we manipulated the weight of the object just after it had been lifted. In this case, the object weight in the lifting phase was different from the weight in the holding phase, thus allowing us to determine what period is most influential in mediating sensorimotor memory. We used a virtual-reality setup to be able to gradually change object weight without changing the physical appearance of the manipulated object and while participants were maintaining contact with the object. Participants lifted objects that were initially light or heavy and, after they had been lifted and were held in the air, their weight could ramp up to heavy or ramp down to light, respectively. We then precisely quantified the force scaling for the next lift to test whether forces were planned based on the weight sensed during lifting or during holding. To compare this behaviour to classic sensorimotor memory experiments, we also included control trials in which the weight of the object did not change and thus was the same during lifting and holding.

## 2 METHODS

### 2.1 Participants

Fifteen right-handed participants (8 females, 21±2 years, age range 18-25 years) took part in the experiment. Participants signed informed consent forms before entering the study. They had no known visual or sensorimotor deficits. The experiment was approved by the local ethical committee of KU Leuven.

### 2.2 Apparatus

The experiment was performed in a virtual reality environment (Figure 1). A 3D-screen (Zalman, 60 Hz screen refresh rate) was reflected on a mirror, under which participants could move their right hand. They inserted their thumb and index finger into the thimbles of two Phantom (Sensable) haptic devices, which provided force feedback to them. The virtual environment showed a patterned background, 4 target positions (yellow marks), the fingertips (red spheres), two start positions (1 for each finger, red-green poles), and a blue rectangular cuboid that was to be lifted. The cuboid measured 5×5 cm in width and 6 cm in height.

**Figure 1.**
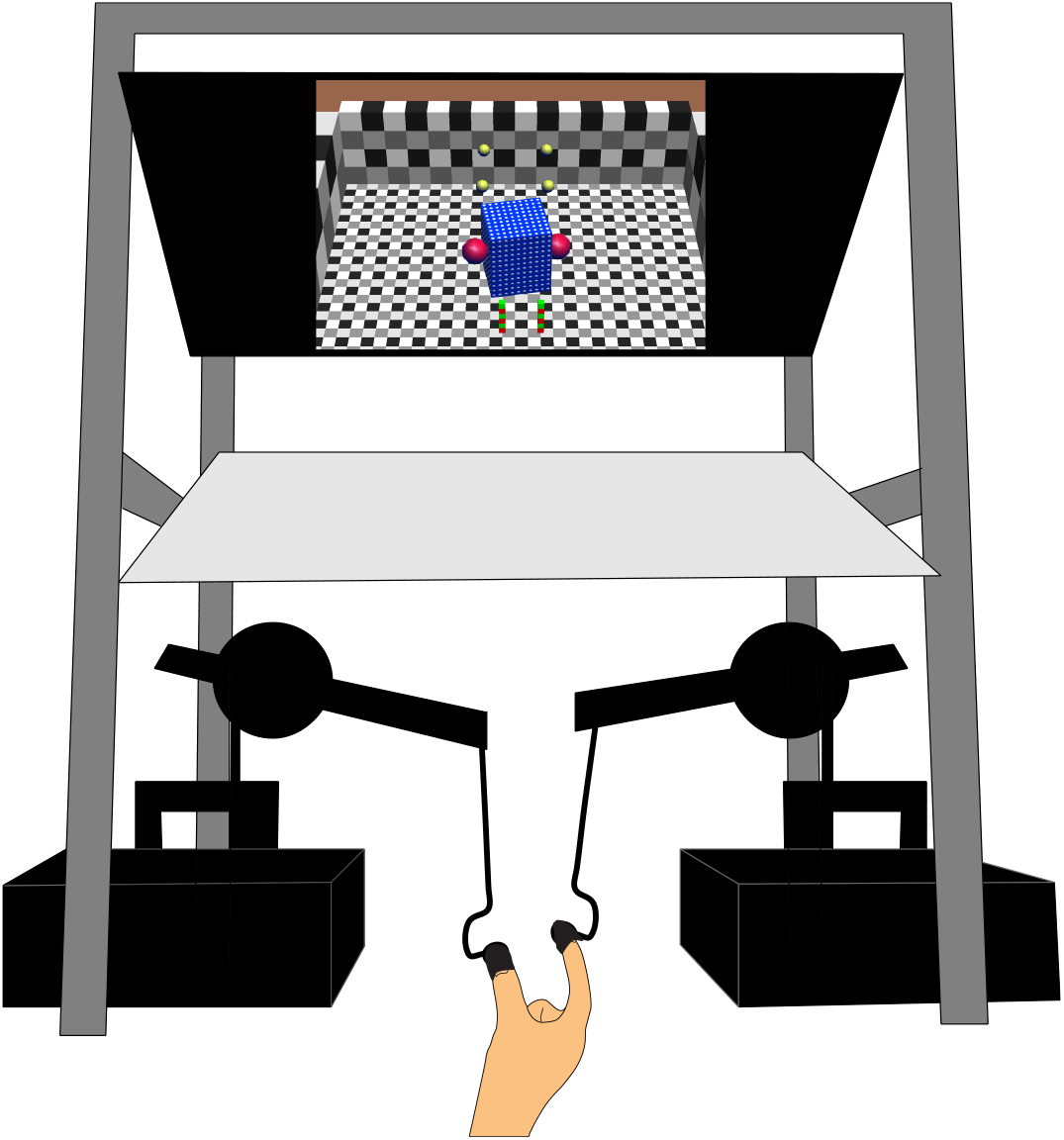
Front view of the virtual reality experimental setup, including a 3D screen, a mirror and two force-feedback robotic devices (Phantom premium 1.5). Participants inserted their thumb and index fingertips into the thimbles of the phantom robots. The virtual environment (only illustrated on the screen) was viewed through the mirror, which projected the scene on the table. The virtual environment showed a blue cuboid, the fingertips (red spheres), target marks (yellow), and fingertip start positions (red-green poles).

A custom-made code written in C++ ensured realistic interactions of the haptic devices with the virtual environment. Horizontal grip forces were simulated as spring-masses. Vertical lift forces were the sum of gravitational, moment and damping forces. Forces and positions in 3 directions of the haptic devices at the fingertips were sampled with a 500 Hz frequency.

### 2.3 Task

Participants were seated in front of the setup with their thumb and index finger inserted the thimbles. They performed practice trials to get familiar with the virtual reality environment. In the experimental trials, they were instructed to grasp the cuboid, lift it until the top corners touched the yellow marks, hold it stable and put it down again. To standardize lifting and holding phases, participants were told to match the timing of their lifting motion to three beeps. The time line of a trial is illustrated in Figure 2. At the start of the trial, the cuboid appeared and they could move their hand from the start positions towards the cuboid, but were instructed not to touch it yet. After the first beep, they could grasp the cuboid and lift it. Participants were required to lift the object from the table up to the target position (yellow marks) within the second beep. Then, they were instructed to hold it stable until the third beep occurred, after which they could return the object to the table. The time between the first and second, and the second and third beep was 1.2 and 1 s, respectively. The first interval was slightly longer to account for reaction times when responding to the first beep.

**Figure 2.**
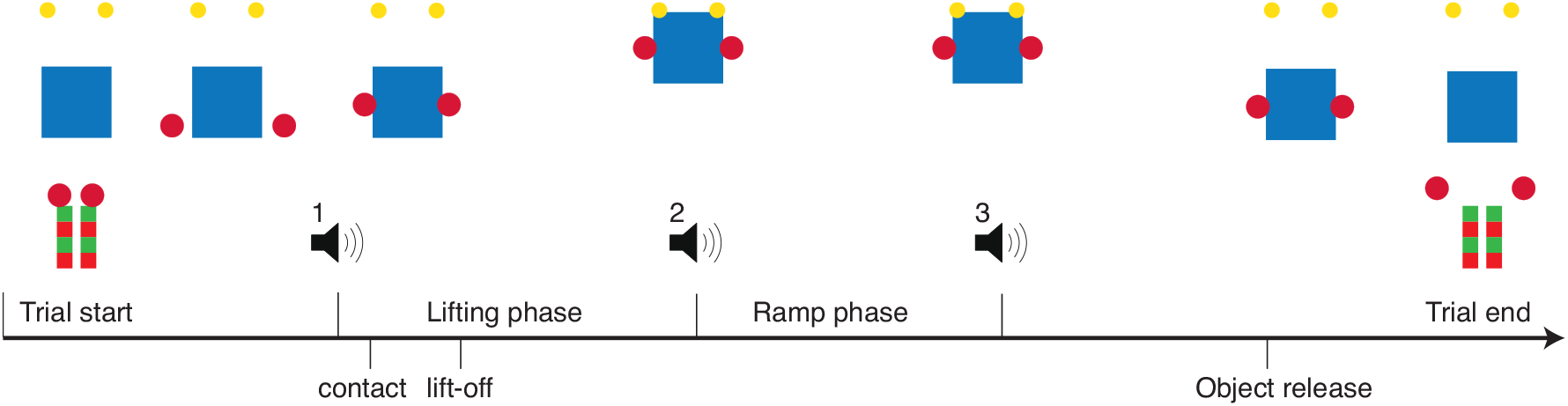
Trial timeline: After the object appeared, participants could move their fingers (red spheres) to the object (blue cuboid) but not touch it yet. After the first beep they could touch it and lift it up to the target level (yellow marks), a position they had to reach within a second beep. Between the second and third beep the object could change in weight (ramp phase). After the third beep participants released the object on the table. The approximate time sequence of object contact, lift-off and object release are indicated on the time line.

The cuboids were of a light (50 g) or heavy (350 g) weight. During the holding phase, i.e. between the second and third beep, the weight of the cuboid could ramp up or down to another weight. A relatively slow ramp was chosen to avoid dropping or moving the object due to sudden weight changes. There were four possible weight combinations in a trial, where the weight was constant (light-light, LL, or heavy-heavy, HH) or changing during hold (light-to-heavy, LH, or heavy-to-light, HL). An example of a LH trial is illustrated in Figure 3. We were interested in the lift after these four possible combinations, which could again be light or heavy. Therefore, there were eight conditions (see Table 1). In the following, a condition is represented as a sequence of light and heavy weights, such as LL-L, where the letters refer to a previous lift, a previous hold and the present object, respectively. Four conditions served as control or ‘no-change’ conditions, which were lifts following trials with a constant weight (LL-L, LL-H, HH-L, and HH-H). The other four conditions were lifts following trials where the weight changed during holding (LH-L, LH-H, HL-L, and HL-H). These were the ‘change’ conditions. Participants performed 15 trials for each condition, resulting in a total of 120 trials that were divided over three sessions. Because the first trial had no preceding lift, it could not be analysed. Therefore, each session consisted of 41 lifts, giving a total of 123 lifts. The whole experiment lasted approximately 90 minutes, including practice and short breaks between the sessions.

**Figure 3.**
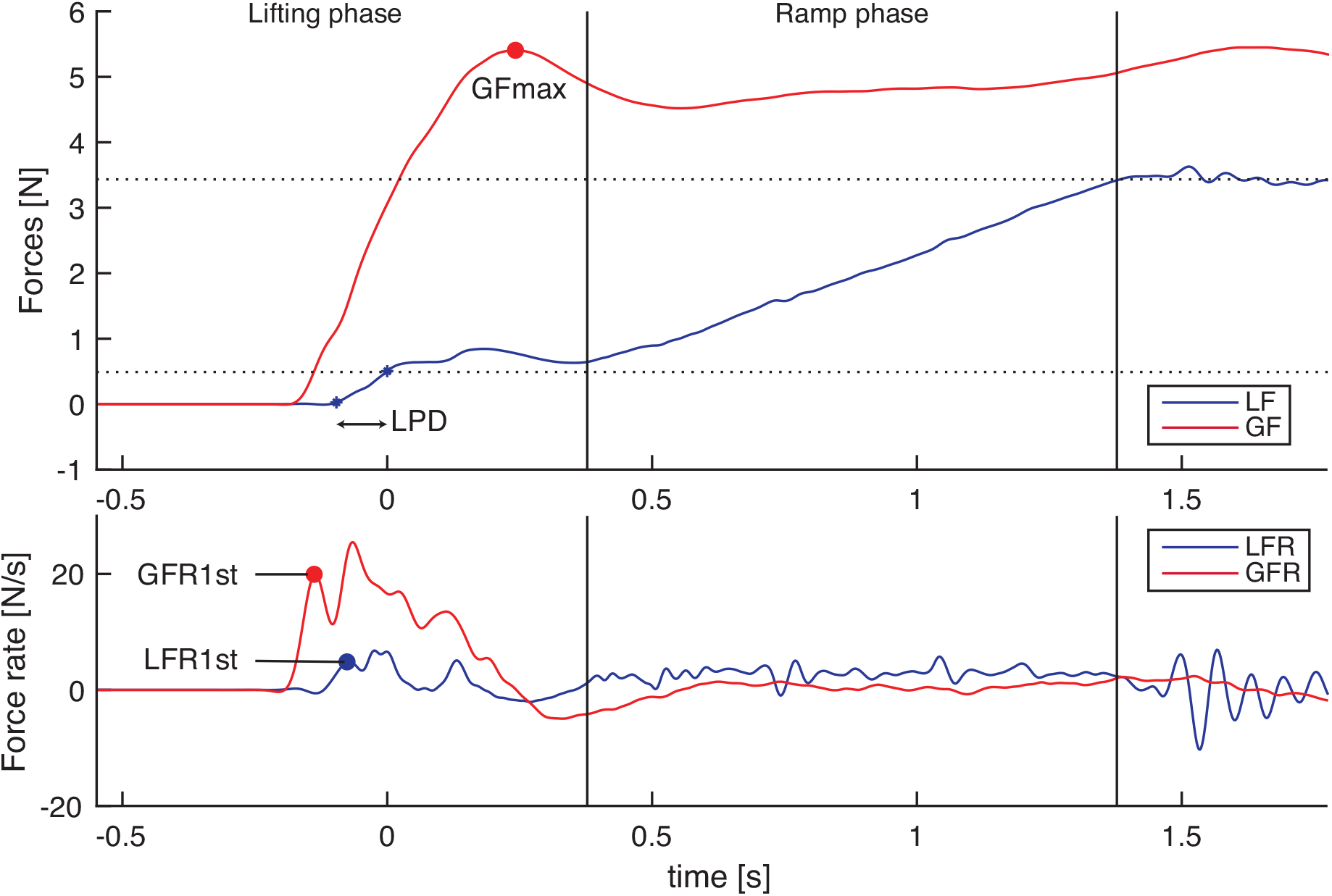
Forces (top) and force rates (bottom) of a single representative trial. Blue and red traces represent load forces (LF) and grip forces (GF), respectively. In this trial, a light object is initially lifted and its weight subsequently ramps up to heavy during holding. Vertical black lines indicate the period during which the object changes weight (between second and third beeps). Horizontal dashed lines indicate the weight of light and heavy objects. The time line is aligned to lift-off. The load force duration (LPD) is the time between LF onset and lift-off. The first peak force rates (GFR1st and LFR1st) and the peak grip force (GFmax) are indicated with filled circles.

**Table 1:** Overview of the conditions. In four conditions, the weight did not change in the previous trial (no-change conditions). In the other four conditions, the weight was different in the lifting and holding phase (change conditions).

|  | Condition | Previous lift | Previous hold | Present object |
| --- | --- | --- | --- | --- |
| No-change conditions | LL-L | Light | Light | Light |
|  | LL-H | Light | Light | Heavy |
|  | HH-L | Heavy | Heavy | Light |
|  | HH-H | Heavy | Heavy | Heavy |
| Change conditions | LH-L | Light | Heavy | Light |
|  | LH-H | Light | Heavy | Heavy |
|  | HL-L | Heavy | Light | Light |
|  | HL-H | Heavy | Light | Heavy |

### 2.4 Analysis

Missing samples (0.002%) were linearly interpolated. Forces were filtered with a second order lowpass Butterworth filter with a cut-off frequency of 15 Hz. To analyse the force scaling in a lift, we calculated the grip forces (GF), defined as the mean of the horizontal forces applied by both fingers, and the load forces (LF), which were the sum of the vertical forces. The grip force rate (GFR) and load force rate (LFR) were the differentiated forces. The onset of GF and LF were based on the force rates and set at the first value after the first beep that was larger than 3 N/s. Lift-off was the first time point the load force overcame the weight of the lifted object. The planning of forces towards object weight is reflected in the value of the first peak of the force rates (Johansson & Westling, 1988a). Therefore, we determined the first peak force rates (GFR1st and LFR1st) after GF onset. To exclude first peaks due to noise or small bumps caused by lightly contacting the object, only first peaks that were at least 10% of the maximum peak rate were considered for analysis. Peak force rates (GFRmax and LFRmax) were calculated as the maximum values between GF onset and 50 ms after lift-off. In addition, we calculated the peak grip force (GFmax) and the load phase duration (LPD). GFmax was the maximum grip force between lift-off and the second beep (i.e. before any weight changes). LPD was the time between LF onset and lift-off. The force parameters of interest are illustrated in Figure 3.

### 2.5 Statistics

The parameters of interest (GFR1st, LFR1st, LPD, GFmax) were averaged for the eight conditions. Outliers (GFR1st: n=1, 0.05%; LFR1st: n=2, 0.11%; LPD: n=7, 0.38%; GFmax: n=0, 0%) that were more than 3 standard deviations away from the mean were removed.

The parameters were analysed with a 2 (previous lift) × 2 (previous hold) × 2 (present object) repeated measures ANOVA. The within factor ‘present object’ was the lifted weight of the object in the current trial. The other two within factors were the object weight of the previously lifted object during the lifting (‘previous lift’) or holding phase (‘previous hold’). Each factor could be light or heavy.

In case of significant interaction effects (*p*<0.05), the separate conditions relevant for the interacting factors were compared with t-tests, using a Bonferroni correction to adjust for multiple comparisons.

To illustrate more clearly the similarity and differences between change and nochange conditions, we performed planned comparisons on the conditions. That is, we compared the ‘change’ conditions to the ‘no-change’ conditions in two ways: 1) with similar weights during lifting, and 2) with similar weights during holding. If force scaling was based on the weight experienced during lifting, change conditions would be more similar to ‘no-change’ conditions with a similar lift and more different from ‘nochange’ conditions with a similar hold.

## 3 RESULTS

Trials with technical errors (4 trials), or where objects were lifted too late (lift-off occurred after the second beep, 39 trials) or too early (contact before the first beep, 20 trials) were removed from analysis. In addition, trials where objects were dropped (9 trials) or released too early (before the third beep, 4 trials) were removed. A total of 76 (4.1%) trials were removed.

A typical force trace profile for the lift of an object whose weight ramps up after the object has been lifted to the target position is shown in Figure 3. The results for GFR1st, LFR1st, GFmax and LPD for each lifting sequence condition are shown in Figure 4, where the first bar stacks (left) are the ‘no-change’ trials without any weight change in the previous trial and the last bar stacks (right) show the ‘change’ trials in which the weight changed during the previous holding phase.

**Figure 4.**
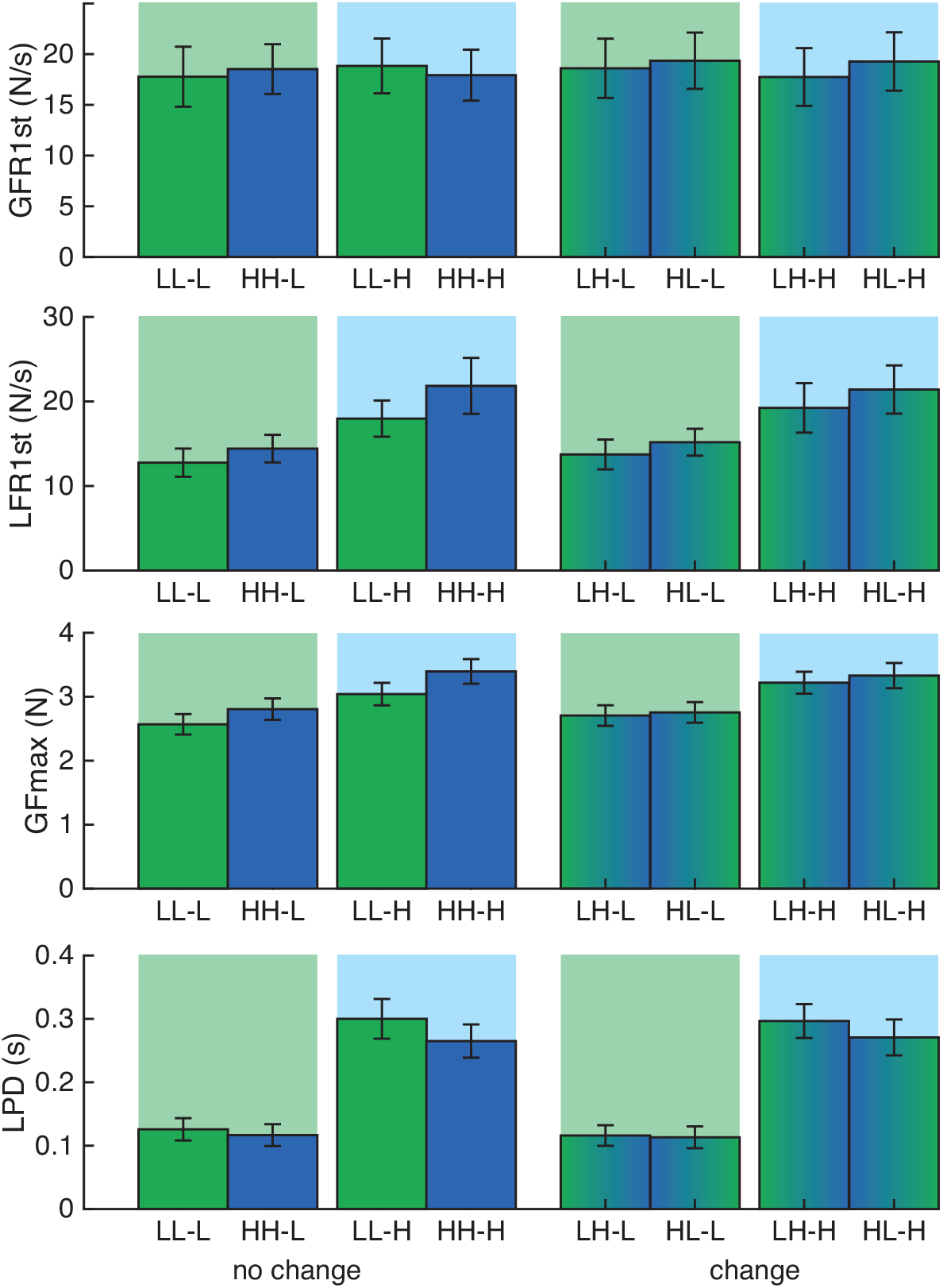
Mean and standard errors for force scaling parameters. From top to bottom: first grip force rate peak (GFR1st), first load force rate peak (LFR1st), peak grip force (GFmax), and load phase duration (LPD). Bars on the left represent no-change conditions (no change in previous lift), bars on the right represent change conditions (weight change in previous lift). Green and blue backgrounds indicate light and heavy lifts of the present object, respectively. Green and blue bar colours indicate previous light and heavy lifts, with the gradient indicating the change in weight during holding. Note the presence of an effect of sensorimotor memory in LFR1st, GFmax and LPD parameters, which only depends on the initial weight (i.e. similar effects for no-change and change conditions).

From previous research, it is expected that when previously lifting a heavy object, force rates for the next lift will be higher than when previously lifting a light object (Johansson & Westling, 1988a). We observed this effect in the load force rates, but not in the grip force rates. For the load force rates, the ANOVA on LFR1st demonstrated significant main effects of previous lift (F(1,14)=20.0, p<0.001, *η*_p_^2^=0.59) and present object (F(1,14)=15.6, p=0.001, *η*_p_^2^=0.53). This showed that if the previous lift was heavy, the first LFR peak was higher than if the previous lift was light. There was no effect of previous hold, suggesting that the weight during this phase had no effect on LFR1st in the next trial. Furthermore, the main effect of present object indicated that the first LFR peak was lower for lifting lighter objects compared to lifting heavier objects. No significant interaction effects were found. The ANOVA on GFR1st showed no significant main or interaction effects. This indicates that, in this experiment, grip force rates were not adjusted towards object weight, neither for the previous or current lift.

In contrast to the grip force rates, the maximum grip forces were affected by previously handled objects. There was a significant effect of previous lift (F(1,14)=34.7, p<0.001, *η*_p_^2^=0.71), previous hold (F(1,14)=9.4, p=0.009, *η*_p_^2^=0.40) and present object (F(1,14)=49.5, p<0.001, *η*_p_^2^=0.78) on GFmax. No significant interaction effects were found, suggesting that these effects were independent. The effect of present object indicated that GFmax was higher for a heavy than for a light object lifted in the present trial. The effect of previous lift showed that the maximum grip force was higher after lifting a heavy compared to a light weight. Surprisingly, also an effect of previous hold was found, suggesting that previously holding a heavy weight resulted in a higher GFmax compared to holding a light weight. However, this is not visible in Figure 3. It is possible that both the effect of previous hold and previous lift were driven by the nochange trials, where the lift and hold weights are the same. In line with this, a separate repeated measures ANOVA on the change trials only, with the factors previous object (light-to-heavy or heavy-to-light) and present object (light or heavy), revealed no effect of previous object (F(1,14)=2.8, p=0.114, *η*_p_^2^=0.17). There was only a main effect of present object (F(1,14)=62.3, p<0.001, *η*_p_^2^=0.82).

For LPD, main effects of previous lift (F(1,14)=17.0, p=0.001, *η*_p_^2^=0.55) and present object (F(1,14)=108.2, p<0.001, *η*_p_^2^=0.89) and an interaction between previous lift × present object (F(1,14)=15.4, p=0.002, *η*_p_^2^=0.52) were found. The main effect of present object indicated that lifting light objects had shorter LPDs than for lifting heavy objects. Post-hoc tests of the interaction of previous lift × present object indicated that this effect was seen both when the previous lift was light (t(14)=–10.7, p<0.001) and heavy (t(14)=9.7, p<0.001). The effect of previous lift showed that LPDs were shorter when the previous lift was heavy compared to previous light lifts. However, this effect was only significant for present heavy objects (t(14)=–4.5, p<0.001) and not present light objects (t(14)=1.6, p=0.124), which explains the interaction effect. There was no effect of previous hold, suggesting that the weight during holding had no effect on the LPD in the next trial.

To test whether change trials were more similar to no-change trials that had the same weight during the lift or to those with the same weight during hold, we compared these conditions separately. In the no-change conditions, the weight of the object did not change during the previous lift. In the change conditions, the object weight in the previous trial was different during the lifting phase than during the holding phase. To test which phase was more influential on the force scaling, we compared the 4 change conditions to the 4 no-change conditions in two ways: 1) we paired the change conditions with no-change conditions based on a similar weight in the lifting phase, and 2) we paired the change conditions with no-change conditions based on a similar holding phase. The comparisons are illustrated in Figure 5. Here, change conditions are presented in green. No-change conditions are presented in blue and red, where in blue the no-change conditions are ordered to match the lifting phase weight of the change conditions, and in red the no-change conditions match the holding phase weight of the change conditions.

**Figure 5.**
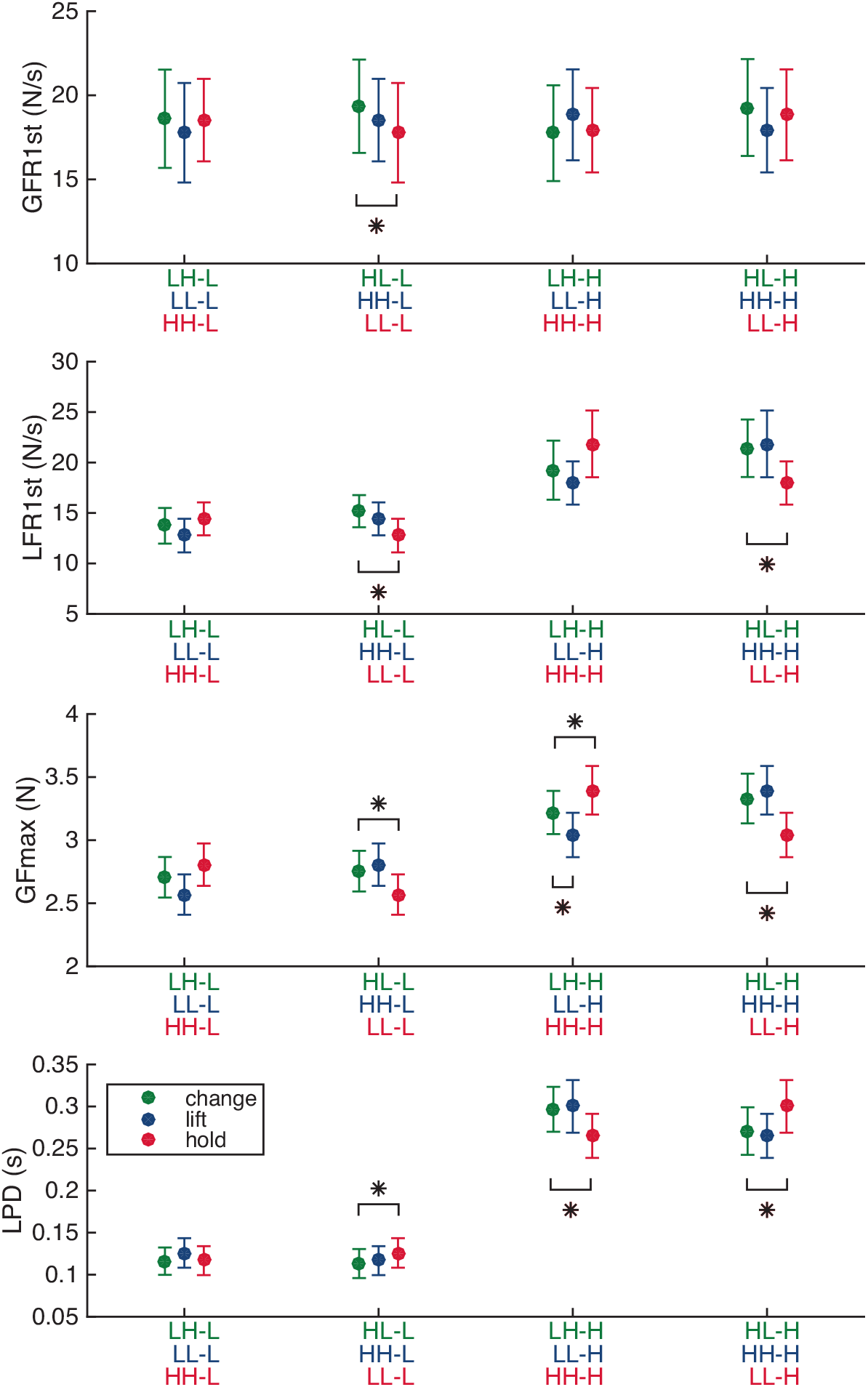
Mean and standard errors for force parameters, comparing change and no change trials. From top to bottom: first grip force rate peak (GFR1st), first load force rate peak (LFR1st), peak grip force (GFmax), and load phase duration (LPD). Green circles indicate change conditions, which are compared to no-change conditions based on similar previous lift weights (blue) or previous hold weights (red). Note that change conditions are more similar to no change conditions when grouped by previous lift rather than by previous hold. *p<0.05.

In comparisons based on the lifting phase, conditions with the same previous lift were compared, irrespective of the holding phase (i.e. LH-L with LL-L, HL-L with HH-L, LH-H with LL-H, and HL-H with HH-H). These are the green and blue data points in Figure 5. Only in the LH-H condition of GFmax, the change condition was different from a no-change condition with a similar lift (t(14)=-4.0, p=0.001). Note that this change condition also differed from the no-change condition with a similar hold weight (see below).

In comparisons based on the hold phase, conditions with the same previous hold were compared irrespective of the lift phase (i.e. LH-L with HH-L, HL-L with LL-L, LH-H with HH-H, and HL-H with LL-H), which are the green and red data points in Figure 5. Significant differences were found for the HL-L condition in GFR1st (t(14)=-2.2, p=0.043). For LFR1st, significant differences were seen in the HL-L (t(14)=-5.1, p<0.001) and HL-H (t(14)=-2.5, p=0.024) condition. In GFmax, the HL-L (t(14)=-4.1, p<0.001), LH-H (t(14)=3.0, p=0.009) and HL-H (t(14)=-5.8, p<0.001) conditions were different from no-change conditions with similar hold. Also for the LPD, the HL-L (t(14)=2.6, p=0.022), LH-H (t(14)=-3.9, p=0.001) and HL-H (t(14)=3.2, p=0.006) conditions differed significantly from the no-change conditions. This indicated that these change conditions were different from the no-change conditions, even though they had similar previous weights in the holding phase. All in all, these results suggest that the behaviour in the change conditions was quite similar to that of no-change conditions with similar previous weights in the lifting phase, but not the holding phase.

## 4 DISCUSSION

The aim of this study was to investigate whether sensorimotor memory for object weight is based on weight experienced during lifting or during holding of an object. Participants lifted light or heavy objects that quickly turned heavy or light, respectively, after the desired lifting height was reached (change trials), We then precisely quantified force scaling on the next lift to determine whether forces were planned based on the weight experienced during the previous lifting or holding phase. Interestingly, we found that forces were higher when previously lifting a heavy object compared to previously lifting a light object, irrespective of the weight experienced during holding. In addition, change trials were more similar to control conditions (no-change trials) with a similar weight experienced during lifting than during holding. Therefore, the main finding of this study is that force scaling is based on a sensorimotor memory of previously manipulated objects that is acquired during lifting, but not holding.

It is surprising that sensorimotor information processed during the lifting phase is the most critical for building up a sensorimotor memory. The lifting phase merely lasts a few hundred ms and is much shorter than the holding phase, thus providing less time to collect sensory information. Strikingly, our findings indicate that the time interval during which information is acquired is more critical than the duration of that interval. Sensorimotor information gathered during the lifting phase might be more important than information during the holding phase because of the several dynamic changes occurring early during lift. Different sensory receptors in the skin respond to dynamic and static stimuli (Johansson & Flanagan, 2009). Similarly, muscle spindles have fibres that are sensitive to either dynamic or static stretch activity (Pearson & Gordon, 2013). The sensitivity of the dynamic and static fibres can be controlled separately which enables setting a higher gain for dynamic sensory inputs during the lifting phase. In such a way, afferent signals from sensors that respond to dynamic changes can be weighted more in the creation of a sensorimotor memory. An argument against this importance of dynamic sensory signals is that, after the holding phase, the object is released, which also represents a dynamic event. Since the object weight during the replacement of the object is the same as that in the holding phase, this suggests that this late dynamic event does not influence sensorimotor memory. Therefore, it is possible that the sensory gain of dynamic inputs is only tuned up during the first phase of object manipulation, possibly to allow for a comparison with expected sensory signals.

During the lifting phase, expected object weight can be compared to actual sensory inputs. Specifically, when a motor command is generated and sent to the muscles, a copy of this command (efference copy) is also generated and can be used to predict the sensorimotor outcome of the action. Actual and predicted sensory signals can be compared to determine whether any adjustments to motor command are necessary and, in turn, update the internal model on which the original motor plan was based (Wolpert & Flanagan, 2001). The lifting phase is the critical time period where the planned forces can be compared to the actual object weight and also adjusted if necessary. Therefore, given that these feedforward sensorimotor loops take place predominantly during the lifting phase, information acquired during this phase might be more critical for building up sensorimotor memory.

Active feedforward control loops are important for building up sensorimotor memories. Previous studies showed that one can better anticipate external force perturbations if they are voluntarily controlled instead of externally induced (Diedrichsen, Verstynen, Hon, Lehman, & Ivry, 2003; Johansson & Westling, 1988b). Additionally, it has been shown that when actively pouring water from a cup, it is subsequently lifted with accurately scaled forces, whereas this is not the case when drinking from the cup with a straw (Nowak & Hermsdörfer, 2003). Such an active component might be crucial because it generates a motor plan with an efference copy which can be used for further predictions.

Another explanation for the importance of the lifting phase for building sensorimotor memory might be that this is the first piece of sensory information received during object manipulation. In other words, the first set of sensory information that is acquired could be more influential and immediately stored for future reference. Sensory signals that are obtained later in handling the object, such as during holding, would have a smaller impact on generating an internal object representation. Indeed, in normal conditions, the object weight should not change between lifting and holding phases, unless, for example, the object is accelerated (e.g. when picking up a cup while driving a car) or filled with liquid (e.g. picking up a glass and pouring water in it). It is therefore plausible that, through daily-life experience, processing of sensorimotor feedback during lifting has been enhanced compared to the holding phase.

It is noteworthy that our effects on the maximum grip force seem at odd with the other force parameters. Specifically, we found both an effect of previous lift and previous hold on the maximum grip force. This seems contradictory, because in the ‘change’ trials, the different experienced weights predict opposite effects on force scaling. For instance, in an LH trial, a low force is predicted based on the weight experienced during lifting, whereas a high force is predicted based on the weight in the holding phase. Still, when comparing the values in the change conditions to the nochange conditions, grip forces were more similar to trials with a similar weight during previous lift than during previous hold. In Figure 4, it seems that the effect of previous lift is stronger, but overall there seems to be no strong effect of previous object experience on the maximum grip force in the change trials. Indeed, when analysed separately, no significant effects of the previous object weight were found, neither during lifting or holding. Therefore, the effect of previous lift and hold might have mainly been driven by the no-change trials. In these trials, both phases have the same weight and predict the same effects. This could suggest a consolidation effect, where experiencing a specific weight for a longer time strengthens the sensorimotor memory. However, such a consolidation effect was not found for the load forces, where force scaling effects seemed similar in change and no-change trials.

Furthermore, there is evidence that the effect of an isometric pinch force on sensorimotor memory is restricted to grip forces (Quaney, et al., 2003), suggesting that grip force may be more influenced by the latest performed force, here experienced during hold, than the load force. It must also be noted that grip forces are not only adjusted to object weight, but also to friction, which might have been more difficult to perceive in the virtual reality setup as no frictional cues are available in the thimbles. Overall, it is plausible that the grip force scaling was more variable. This is also apparent in the absence of any scaling effects in the grip force rates in both the nochange and change conditions.

In conclusion, we show that sensorimotor memories underlying object manipulation are based on information that is acquired during lifting, but not holding. The feedforward comparison between expected and actual sensory inputs taking place within this particular time period could be more critical than information acquired during static holding for the generation of internal object representation.

## ACKNOWLEDGEMENTS

VVP is funded by a Fonds Wetenschappelijk Onderzoek grant [Post-doctoral fellowship, FWO, Belgium, 12X7118N]. MD is funded by a Fonds Wetenschappelijk Onderzoek grant [FWO Odysseus, Belgium FWO: G/0C51/13N]. The authors would like to thank Dr. Atsuo Nuruki and Dr. Raz Leib for their help in designing the virtual environment.

